# Isolation of functional lysosomes from skeletal muscle

**DOI:** 10.1101/2025.09.09.675181

**Authors:** Thulasi Mahendran, Anastasiya Kuznyetsova, Neushaw Moradi, David A. Hood

## Abstract

Lysosomes are membrane-bound organelles responsible for the degradation of damaged or dysfunctional cellular components, including mitochondria. Their acidic internal environment and the presence of an array of hydrolytic enzymes facilitate the efficient breakdown of macromolecules such as proteins, lipids, and nucleic acids. Mitochondria play a critical role in maintaining skeletal muscle homeostasis to meet the energy demands under physiological and pathological conditions. Mitochondrial quality control within skeletal muscle during processes such as exercise, disuse, and injury is regulated by mitophagy, where dysfunctional mitochondria are targeted for lysosomal degradation. The limited understanding of quality control mechanisms in skeletal muscle necessitates the need for isolating intact lysosomes to assess organelle integrity and the degradative functions of hydrolytic enzymes. Although several methods exist for lysosome isolation, the complex structure of skeletal muscle makes it challenging to obtain relatively pure and functional lysosomes due to the high abundance of contractile proteins. Here we describe a method to isolate functional lysosomes from small amounts of mouse skeletal muscle tissue, preserving membrane integrity. We also describe functional assays that allow direct evaluation of lysosomal enzymatic activity and we provide data indicating reduced lysosomal degradative activity in lysosomes from aging muscle. We hope that this protocol provides a valuable tool to advance our understanding of lysosomal biology in skeletal muscle, supporting investigations into lysosome-related dysfunction in aging, disease, and exercise adaptations.

**New and Noteworthy:** Lysosomes within skeletal muscle function to degrade dysfunctional debris and initiate retrograde signaling pathways. We developed a method to isolate purified lysosomal fractions using a small portion of skeletal muscle, eliminating the need for density gradients or lysosome-modifying agents, ensuring high lysosomal purity without compromising structure or function. By enabling functional analysis via acid phosphatase, cathepsin-B activity, and calcium release, this approach offers a powerful tool to study lysosomal roles in muscle physiology, disease, and exercise.

**Graphical Abstract:** 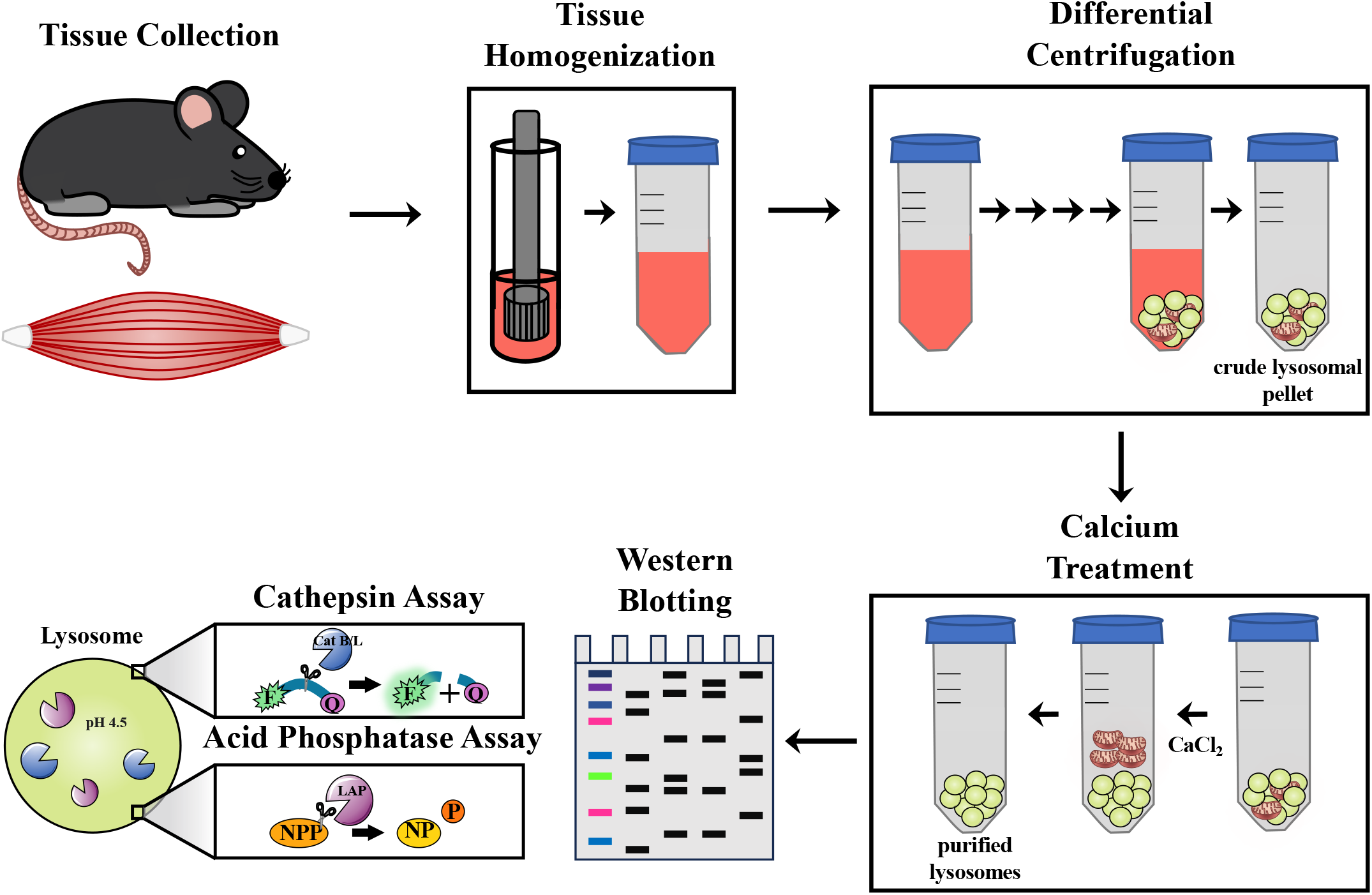

## Introduction

The regulation of skeletal muscle health involves the coordinated activity of cellular organelles and their capability to adapt to various physiological conditions (1). Lysosomes are membrane-bound organelles designed for the degradation of dysfunctional cargo. They are characterized by an acidic interior and an extensive repertoire of hydrolytic enzymes that operate optimally at a low pH to break down molecules. This acidic microenvironment facilitates the degradation of macromolecules delivered via endocytosis, phagocytosis, and autophagy, ensuring the recycling of essential nutrients and the clearance of cellular debris (2). Mitochondria are organelles which produce energy for muscle contraction and can be increased in both content and function in skeletal muscle by the adoption of a regular exercise program (3). In contrast, mitochondrial content and function can be reduced within the cell by extended periods of muscle disuse, thereby altering cellular energy metabolism (4). Dysfunctional mitochondria may contribute to this decline in energy metabolism when pathological but can also trigger adaptive signaling pathways that can act as intentional mechanisms to maintain homeostasis (5). The process of mitophagy mediates the removal of dysfunctional mitochondria via their delivery to lysosomes for degradation. The terminal step of mitophagy involves the fusion of autophagosomes containing defective mitochondria with lysosomes for degradation and recycling (6). This function is vital for the clearance of debris which could otherwise aggregate the existing mitochondrial network and have pathological consequences (7). Since lysosomes play a critical role in maintaining mitochondrial homeostasis via mitophagy, poorly functioning lysosomes and hydrolases within will affect the preservation of skeletal muscle health (7). Evidence for lysosomal dysfunction is evident from the presence of lipofuscin viewed using electron microscopy techniques in denervated (8) and aging (9) muscle. Lysosomes are also involved in retrograde signaling pathways where they serve as hubs for mediating cellular signaling, such as the mechanistic target of rapamycin complex 1 (mTORC1). Lysosomes also release calcium and promote the activation of the MiTF/TFE transcription factor family of proteins such as TFEB and TFE3 that act in the nucleus to orchestrate lysosomal biogenesis in response to nutrient availability, stress stimuli, and growth factors (8). While these functions and signaling pathways have been appreciated in many cell types, little information exists with respect to skeletal muscle.

Our previous work has shown that chronic exercise leads to improved mitochondrial function and content and can induce coincident increases in lysosomal proteins as well (9-11). This suggests that circumstances of reduced lysosomal function could potentially be rescued by exercise through lysosomal biogenesis and intra-organelle adaptations. However, the mechanisms leading to lysosome synthesis and functional changes are unclear. Importantly, it should be noted that changes in the expression of lysosomal markers does not necessarily translate or coincide with increases in the function of the entire organelle. This can best be assessed using functional assays on isolated lysosomal fractions. While a number of methods have arisen to isolate lysosomes (12-14), few of these (15, 16) have been applied to skeletal muscle. Muscle is relatively difficult to study in this respect because of the robustness of the tissue and the abundance of contractile proteins which dominate the cellular landscape making it difficult to isolate organelles. A procedure for the isolation of lysosomes would be a very useful technique to address the knowledge gap regarding putative lysosome functional defects in disease conditions, as well as with aging and/or the adaptation to regular exercise. Thus, the objective of this paper was to develop a technique for the isolation of lysosomes from relatively small amounts of skeletal muscle that can be used for functional assays.

## Materials and Methods

### Animals and tissue extraction

C57BL/6J mice were obtained from Jackson Laboratories. Animals were bred in accordance with the guidelines of the York University Animal Care Committee. Tissue extraction was performed under anesthesia with isoflurane. Hindlimb skeletal muscle was exposed after surgically removing skin and fascia. The tibialis anterior was located lateral to the tibia and extracted via the muscle closest to the origin and the distal tendon. To locate and expose the gastrocnemius and soleus, the calcaneal tendon at the distal insertion was severed. The soleus, located deep to the gastrocnemius, was excised via its origin and insertion. The gastrocnemius was extracted via the origin closest to the femur. The quadriceps was extracted by positioning the hindlimb at a 90° angle and cutting along the femur towards the origin. Fresh skeletal muscle samples were utilized for isolation of lysosomes. After skeletal muscle collection, mice were euthanized via removal of the heart.

### Lysosome isolation

The overall scheme for the isolation of lysosomes is illustrated in Fig. 1. Fresh skeletal muscle tissue (1.0 g) was isolated and washed twice in ice-cold PBS. The tissue was then placed in a pre-chilled glass Petri dish and minced on ice using sharp scissors. The minced tissue was weighed and transferred to a 50 mL conical tube and 5X volume of Lysosome isolation buffer (LIB) comprising 250 mM sucrose, 50 mM Tris–HCl pH 7.4, 5 mM MgCl_2_ was added. The tissue suspension was then homogenized for 20 sec using a tissue homogenizer (T25 digital ULTRA-TURRAX®) at 9,200 rpm. This step was repeated two more times. The homogenate was decanted into a 15 ml centrifuge tube and maintained on ice until placed in the centrifuge. The minced muscle suspension was centrifuged at 1,000 g for 10 min at 4 C and the supernatant was transferred to a new 15 ml conical tube. The pellet was resuspended in 2X volume of LIB and homogenized once at 9,200 rpm for 20 sec. The sample was centrifuged at 1,000 g for 10 min and the supernatant was transferred to the previous tube. The pellet containing nuclei and any tissue debris was discarded. The collected post-nuclear supernatant (PNS) was centrifuged at 3,000 g for 13 min at 4 °C. The pellet (HMF1) was discarded, and the supernatant was recentrifuged at 3,000g for 13 min to remove any remaining heavy mitochondria. Following centrifugation, the supernatant was transferred to a new 15 ml tube and the pellet discarded. The supernatant (HMFS2) was centrifuged at 9000g for 10 min and the pellet containing light mitochondrial fraction (LMF1) was discarded. This step was repeated once more at 9,000g for 10 min to further remove light mitochondria (LMF2) from the supernatant (LMFS2). Purified LMFS2 was then centrifuged at a higher speed to remove the cytosolic fraction (Cyto) and enrich the lysosomes in the form of a pellet. The pellet was resuspended in 400 µl of LIB containing 8 mM CaCl_2_. The suspension was rocked on an orbital shaker at 4 °C for 30 min followed by centrifugation at 5,000 g for 15 min. The pellet containing lysosomes was then washed 3 times at 5,000 g for 10 min in 500 μl LIB for further purification and the final pellet was resuspended in 40 to 60 μl of LIB (**Fig. 1**).

**Figure 1.**
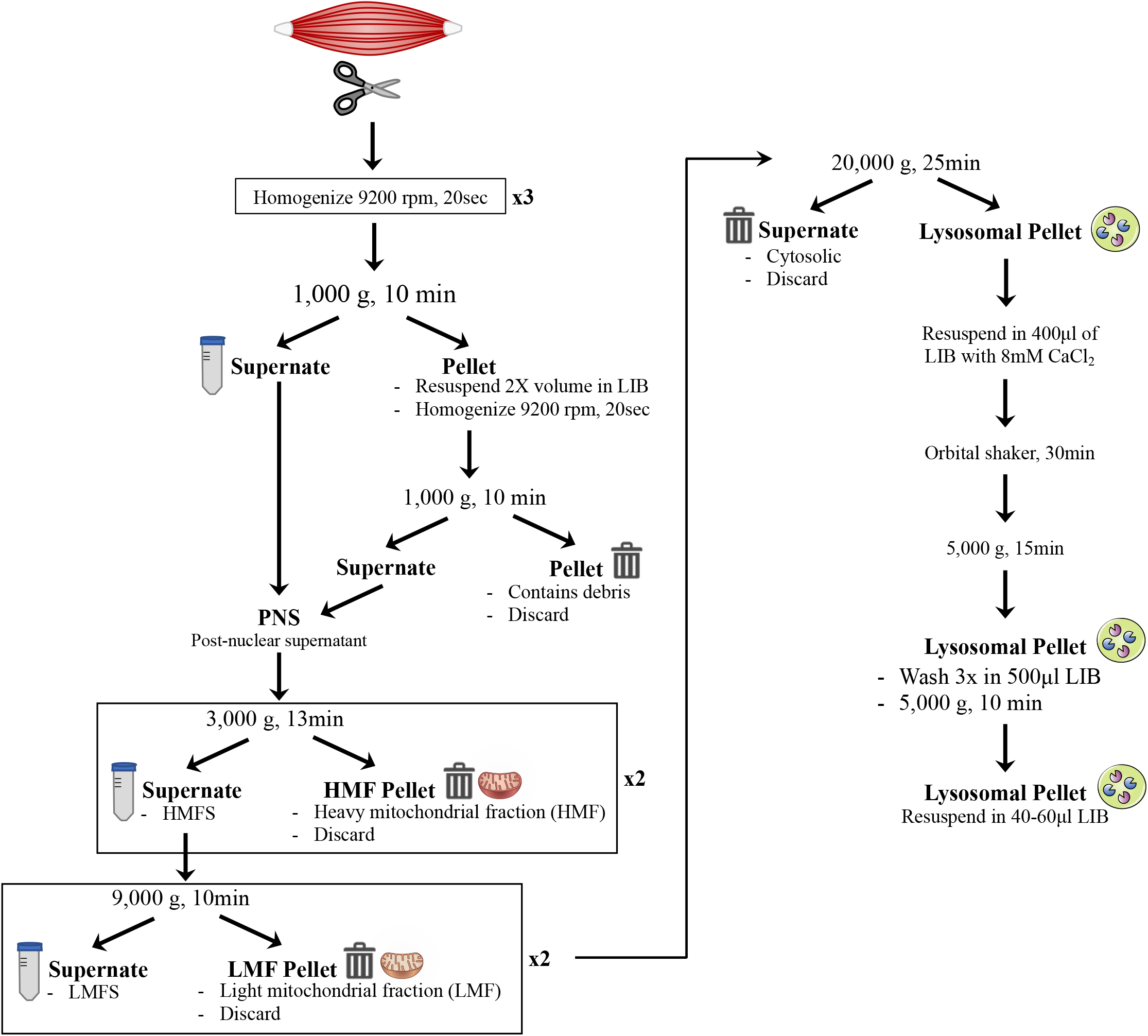
Schematic steps of lysosome isolation from fresh minced skeletal muscle, described in detail in Methods: Lysosome Isolation. LIB, lysosome isolation buffer; HMF, heavy mitochondrial fraction; LMF, light mitochondrial fraction.

### Cathepsin activity assay

The fluorogenic peptide substrate Z-Leu-Arg-AMC (Cat. No. ES008) was purchased from R&D Systems. A 10 µM concentration of the substrate was incubated with 10 µg of lysosomal suspension (protein-based) in MES buffer (pH 5.0) in a 100 µL reaction volume at RT for 10 minutes. The reaction mixtures were then transferred to a 96-well plate, and enzymatic activity was measured via fluorescence, with excitation at 380 nm and emission at 460 nm over a 45 min period. The fluorescence emission of each sample was normalized by subtracting the background fluorescence generated by the peptide substrate in MES buffer alone. The gradient of the initial linear region of the fluorescence curve was used to calculate the enzymatic activity rate of Cathepsins B/L in the lysosomal extract. A higher fluorescence signal corresponded to increased enzymatic activity, reflecting the enrichment of active Cathepsins B/L in the lysosomal fractions.

### Acid phosphatase assay

Substrate Solution containing 4-nitrophenyl phosphate was made by dissolving a substrate tablet in 2.5 ml of citrate buffer. The substrate solution was prepared freshly each time and equilibrated to 37 °C prior to use. The reaction mixtures were prepared by mixing 50 µL of Substrate Solution, 50 µL of the lysosomal suspension in citrate buffer. The blank was made by mixing 50 µL of substrate solution and 50 µL of citrate buffer without lysosomal suspension. The reactions were mixed thoroughly by pipetting and the tubes were incubated at 37 °C for 30 min. Reactions were then stopped by adding 100 µL of stop solution and transferred to a 96-well plate. Absorbance was measured at 405 nm using a plate reader and the absorbance readings were normalized by subtracting the reagent blank values to account for background hydrolysis of the substrate.

### Immunoblotting

Protein concentrations of samples were determined using the Bradford assay. Samples were loaded onto 10-18% SDS-PAGE gels and separated by electrophoresis at 120V. Following separation, gels were transferred onto nitrocellulose membrane (Bio-Rad). Membranes were then stained with Ponceau red and cut according to where protein of interest falls based on its molecular weight. The cut membranes were washed with 1xTBS-T (tris-buffered saline – Tween) for 5 minutes, then blocked with 5% milk in 1xTBS-T or 5% BSA (bovine serum albumin) in 1xTBS-T for 1 hour at room temperature. Membranes were then laid out on a flat surface and incubated overnight at 4°C with their appropriate primary antibody (Table 1). The next morning, membranes were washed 3 times in 5-minute intervals with 1xTBS-T. After washing, the membranes were incubated for 1 hour at room temperature with their appropriate secondary antibody. The membranes were again washed 3 times in 5-minute intervals with 1xTBS-T before beginning imaging of the membrane blots. Membranes were imaged using an ECL kit (170-5061; Bio-Rad) on the Invitrogen iBright 1500 imager (ThermoFisher). The blots were quantified using ImageJ software (ver. 1.53) and GraphPad Prism Software (ver. 8.2.1).

**Table 1.** List of antibodies used for immunoblotting.

| Antibody | Concentration | Company | Catalog Number |
| --- | --- | --- | --- |
| Catalase | 1:1000 | Cell Signaling | 14097S |
| Cathepsin B | 1:1000 | Cell Signaling | 31718S |
| COX IV | 1:1000 | Abcam | AB14714 |
| Lamp1 | 1:1000 | Abcam | AB24170 |
| LC3 | 1:1000 | Cell Signaling | 4108S |
| mTORC1 | 1:1000 | Cell Signaling | 2972S |
| Rab5 | 1:1000 | ABclonal | A23955 |
| Rab7 | 1:50,000 | ABclonal | A12308 |
| α-tubulin | 1:10,000 | Millipore-Sigma | CP06 |
| HRP-linked anti-mouse | - | Cell Signaling | 7076S |
| HRP-linked anti-rabbit | - | Cell Signaling | 7074S |

### Electron Microscopy

Isolated lysosome pellets for TEM imaging were fixed in 2% glutaraldehyde in 20 mM phosphate buffer containing 0.25M sucrose for 2 hours. Pellets were then rinsed in buffer, followed by post-fixation in 1% osmium tetroxide in buffer for 90 min and dehydration in a graded ethanol series (50%,70%,90%, and 100%, 20 minutes each step). Dehydrated pellets were subjected to two propylene oxide changes for 30 min and embedded in Quetol-Spurr resin. Blocks were cured overnight in the oven at 60 °C. Sections 50 nm thick were cut on a Leica EM UC7 ultramicrotome, and post-stained with 2% uranyl acetate and 3% lead citrate for 20 minutes each and washed for 5 minutes after each staining. Sections were air dried at room temperature before viewing under TEM.

### Flow cytometry

Isolated lysosomes were stained in 100 nM Lyso Tracker Green (LTG, Thermo Fisher Scientific) for 45 min at 37 °C while shaking. The sample was then centrifuged at 5000 g for 10 min at 4 C. The pellet was washed and resuspended in 200 μl PBS. Analysis was performed using a Beckman CytoFLEX Flow Cytometer. LTG staining was detected using a Fluorescein Isothiocyanate (FITC) filter with a 488 nm laser. The sample volume was set to 10 μL, and the flow rate was limited to a maximum of 3000 events/sec.

### Calcium Kinetics Assay

The protein concentration of each lysosomal preparation was assessed using the Bradford assay. Protein-matched isolated lysosomes were then stained with 3 μM of FLUO-4 AM (Invitrogen; Cat No. F14201) in lysosomal isolation buffer (LIB) and incubated at 37°C for 30 minutes. Following this, samples were centrifuged at 21 000 x g for 10 minutes at 4°C. The supernatant fraction was removed and replaced with an arbitrary volume of new LIB to the remaining pellet for a de-esterification process. The samples were subsequently incubated at 37°C for 30 minutes and then were centrifuged again at 21 000 x g for 2 minutes at 4°C for the removal of any LIB that contained any remnant extra-lysosomal FLUO-4 dye. The remaining pellet for each sample was volumed up to 100 μL to be sufficiently dispensed into a black, glass-bottom 96-well plate (CellVis/Cedarlane; Cat No. P96-1.5H-N). Using the CYTATION5 plate reader (BioTek), each dispenser was primed with a prepared volume of the TRPML1 synthetic agonist ML-SA1 (Sigma; Cat No. SML0627-25MG) or DMSO, with LIB as the solvent. The final concentration of TRPML1 (and DMSO) was 10 μM. The temperature of the plate reader was set to 37°C. During read time, a basal signal was collected for each sample for 20 seconds 50 millisecond intervals. Following this, 10 μL of DMSO or ML-SA1 was dispensed into each well, followed by a kinetic read of 60 seconds at 50 millisecond intervals. The FLUO-4 AM fluorescent wavelengths were set to 485/20 nm excitation, and 528/20 nm emission.

## Results

The purity of the lysosomes can be determined by the relative enrichment of lysosomes and by the absence of contaminants such as the major sub-cellular organelles and cytoplasm. We measured the levels of the lysosomal marker LAMP1, the cytosolic marker α-Tubulin, the peroxisomal marker catalase and the mitochondrial marker COX4 using western blot analysis to determine the enrichment and purity of the lysosomal fraction (**Figs. 2A, 2B**). Following lysosome isolation using the described protocol (**Fig. 1**), an increase in the intensity of lysosomal markers, such as mature Cathepsin B (6.8-fold) and LAMP1 (11.8-fold), was observed in the lysosomal fraction compared to the tissue homogenate, while the levels of Tubulin and catalase were significantly reduced (0.13-fold and 0.21-fold, respectively) under the same exposure conditions. This reduction indicated that the isolated fraction was free of cytoplasmic and peroxisomal components. However, a significant amount of COX4 (4.2-fold) was detected in the lysosome-enriched fraction, at levels comparable to those in the heavy and light mitochondrial fractions. This observation indicates contamination of the isolated lysosomal fraction with mitochondrial components. These initial observations required us improve the purification method to achieve a fraction with a relatively higher degree of purity without any mitochondrial contamination.

**Figure 2.**
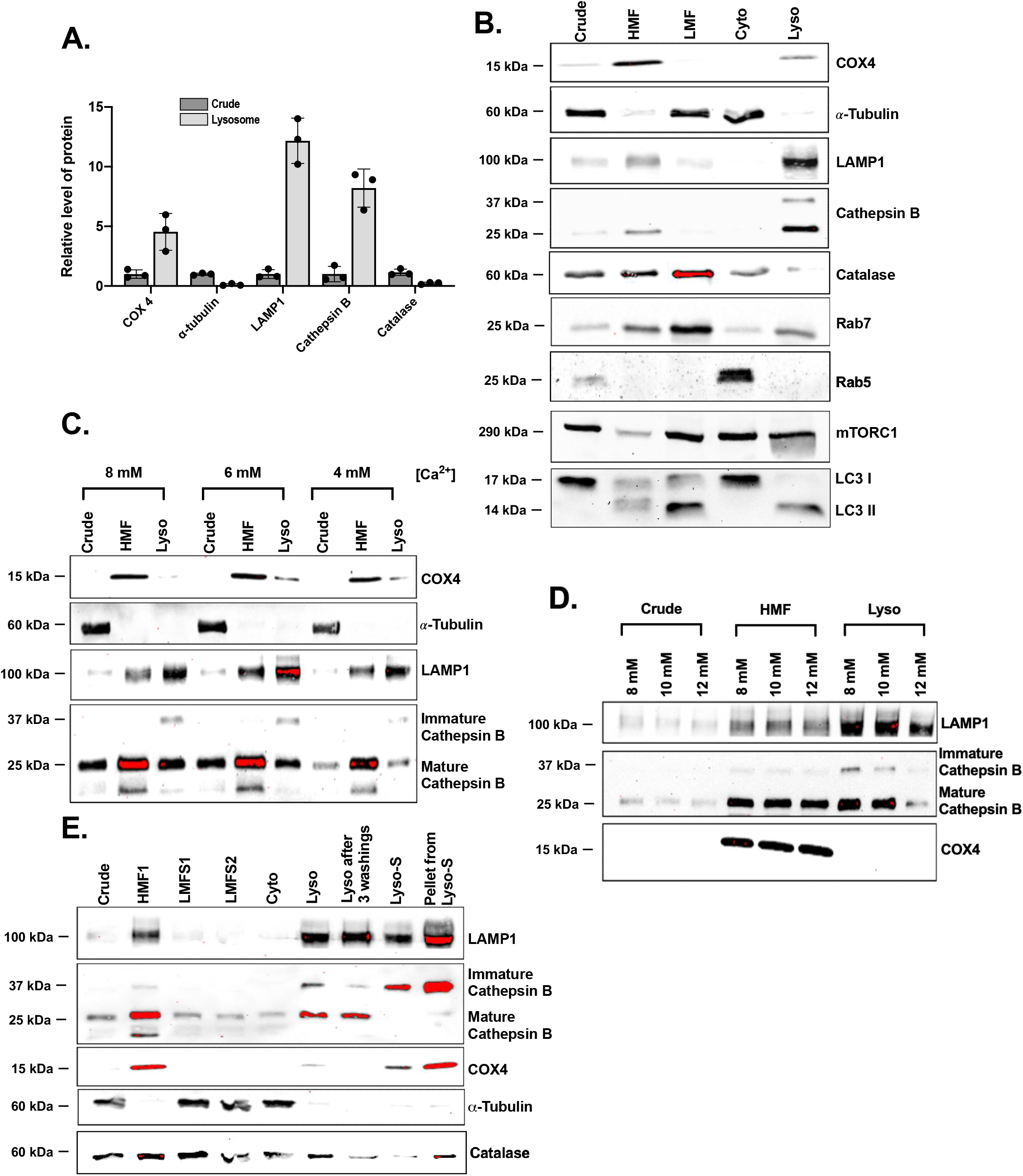
Western blot analysis for different subcellular organelle markers during lysosomal isolation and fold enrichment of LAMP1, Cathepsin B, COX4, α-Tubulin, Catalase, Rab7, Rab5, mTORC1, LC3 I, and LC3 II. *A, B*: The representative blot used to determine the fold changes (FCs). *C*, D: Western blot demonstrating the removal of contaminating mitochondria by varying Ca^2+^ concentration during lysosomal isolation. *E*: Western blot analysis of fractions collected during lysosome isolation after Ca^2+^ and centrifugation optimizations. HMF, heavy mitochondrial fraction; LMF, light mitochondrial fraction; LMFS, light mitochondrial fraction supernatant; S, supernatant.

Based on our initial protocol, the heavy mitochondrial fraction was isolated by centrifugation at 3,000 g for 10 min, followed by isolation of the light mitochondrial fraction at 9,000 g for 10 min. To facilitate the removal of mitochondria we modified these steps by adjusting the centrifugal speed and duration, which led to significant improvements in the resulting blots. Additionally, washing the lysosomal pellet three more times helped achieve a much purer fraction free from mitochondrial contamination. The addition of Ca^2+^ leads to mitochondrial swelling and reduces the density of mitochondria, which in turn aids in the removal of any contaminating mitochondria from lysosomes, a technique that has been previously used during density gradient isolation of lysosomes from rat liver (12). During the optimization process, we also varied the concentration of calcium (Ca^2+^) used to remove residual mitochondria, ranging from 4 mM to 12 mM (**Fig. 2C and 2D**). The lysosomal fraction obtained with a Ca^2+^ concentration of 8 mM showed the highest LAMP1/COX4 ratio, indicating minimal mitochondrial contamination. Concentrations below 8 mM resulted in higher contamination (**Fig. 2C**), while concentrations above 8 mM, although effective in fully removing mitochondria, led to a decrease in lysosomal yield (**Fig. 2D**). These modifications significantly improved the purity of the lysosomal fraction, providing a reliable protocol for isolating lysosomes with minimal mitochondrial contamination (**Fig. 2E**).

In addition to the above-mentioned organelle markers, we also probed for Rab5 to detect the presence of early endosomes. The absence of Rab5 confirmed the enrichment of mature lysosomes with minimal early endosomal contamination (**Fig. 2B**). At the same time, we noticed the presence of Rab7 in the isolated lysosomal fraction (**Fig. 2B**). This is in support of the fact that Rab7 is associated with late endocytic organelles, predominantly early lysosomes with degradative capacity (17, 18). Apart from these markers, we also examined LC3-II, which is a crucial marker for autophagy (19). Our blots show the presence of LC3-II in the lysosomal fraction, suggesting the presence of autolysosomes in our lysosomal pool. This indicates that the lysosomes are functionally active and engaged in autophagic processing. Moreover, the detection of LC3-II supports the fact that our fractions represent a heterogeneous pool of lysosomes containing not only terminal lysosomes but also autolysosomes, resembling the physiological lysosomal population. In addition, since lysosomes serve as signaling hubs particularly for nutrients and energy homeostasis, it was essential to confirm that the signaling molecules associated with lysosomes remain intact during sample processing. mTORC1, a key nutrient-sensing kinase, is known to be recruited to the lysosomal surface in response to nutrient availability (24). Our western blot analysis confirmed the presence of mTORC1 in the lysosome-enriched fraction, supporting the retention of lysosome-associated signaling components during isolation (**Fig. 2B**).

We also determined whether the isolation protocol yielded intact lysosomes and preserved the molecular integrity of the lysosomal membrane by performing TEM imaging. This technique allows the detection of lysosomes as single membrane-enclosed organelles within a size range of 50 – 600 nm (20-22). The TEM image (**Fig. 3A**) confirmed that the lysosomes isolated were intact, enclosed by a single unit membrane, and within the nanometer size range. Though most of them appeared circular, we observed variations in the size and morphology of the lysosomes within the isolated pool, indicating that they are at different stages of the lysosome cycle (23). In addition, our images also confirmed the absence of mitochondrial contamination. To corroborate TEM data and validate the intactness of enriched lysosomes, we incubated them with pH-dependent lysosomal fluorescent dye lysotracker green (100 nM). Flow cytometry analysis confirmed the retention of the LTG within the lysosomes (∼68% LTG+), suggesting that the isolated lysosomes were intact (**Fig. 3B and 3C**).

**Figure 3.**
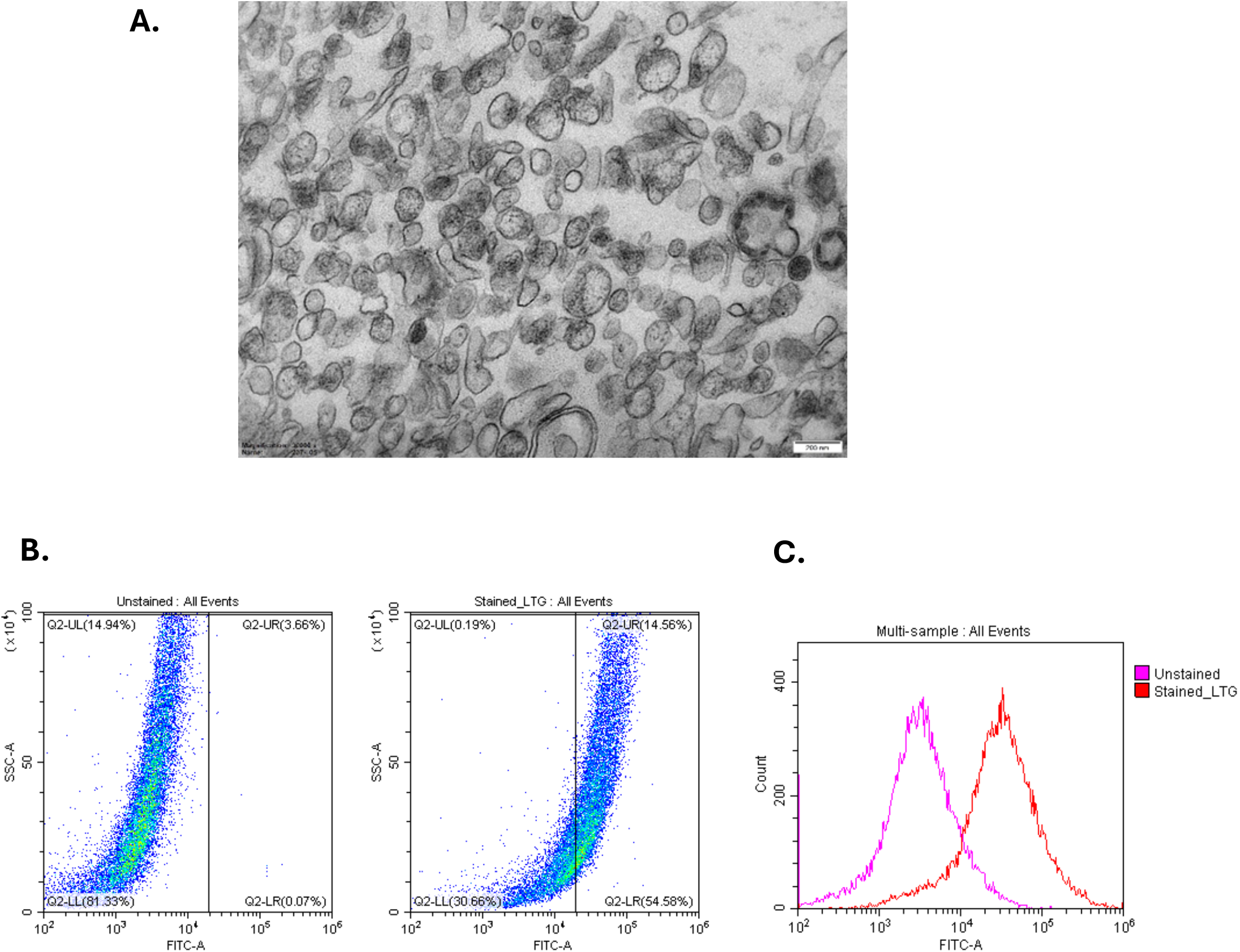
*A*: Transmission electron micrograph representing the pool of lysosomes in the purified lysosomal fraction (Scale bar: 200 nm and Magnification: 30,000x). *B*: Representative flow cytometry scatter plots for unstained and LTG (100 nM) stained lysosomes. *C*: Representative flow cytometry histogram showing the overlay of the unstained and stained lysosomes.

To evaluate the functionality of intact lysosomes, we employed three distinct methods: 1) measuring the activity of lysosomal cathepsins B/L, 2) assessing the activity of lysosomal acid phosphatase (LAP), and 3) quantifying lysosomal calcium release. Lysosomal acid phosphatase is initially synthesized as a transmembrane protein and transported to lysosomes, where it undergoes proteolytic cleavage to form its mature, soluble form (25). LAP activity was assessed using 4-nitrophenyl phosphate as the substrate, monitoring the increase in absorbance at 405 nm due to the formation of the yellow-colored reaction product, p-nitrophenol. The assay was performed with varying concentrations of the isolated lysosomes, revealing a linear increase in activity (**Fig. 4C**), which confirmed the integrity of the isolated lysosomes. Additionally, activity was measured over different incubation periods at 37 °C, showing a time-dependent increase (**Fig. 4D**). Together, these results demonstrate that the isolated lysosomes are functionally active. In addition, we also observed a ∼15-fold increase in LAP activity in the purified lysosomes compared to the crude fraction. The isolated lysosomal fractions were further tested for Cathepsin B/L activity using a fluorescence-based reporter assay. Lysosomal cathepsin proteases are initially synthesized as inactive zymogens, transported to endosomes for maturation, and subsequently transferred to lysosomes where they become fully active (26). The fluorogenic substrate Z-Arg-Arg-AMC used in this study can be cleaved by Cathepsins B and L under acidic pH resulting in an emission at 460 nm (**Fig. 4A**). The assay demonstrated that these cathepsins were active in the isolated lysosomal fractions as we observed an increase in the fluorescence emission in a time course of 45 min due to the cleavage of the substrate (**Fig. 4B**). Further, we compared the activity of these enzymes between the whole-muscle homogenate and the purified lysosomal fraction, where we observed a ∼5-fold activity enrichment in the latter (**Fig. 4B**).

**Figure 4.**
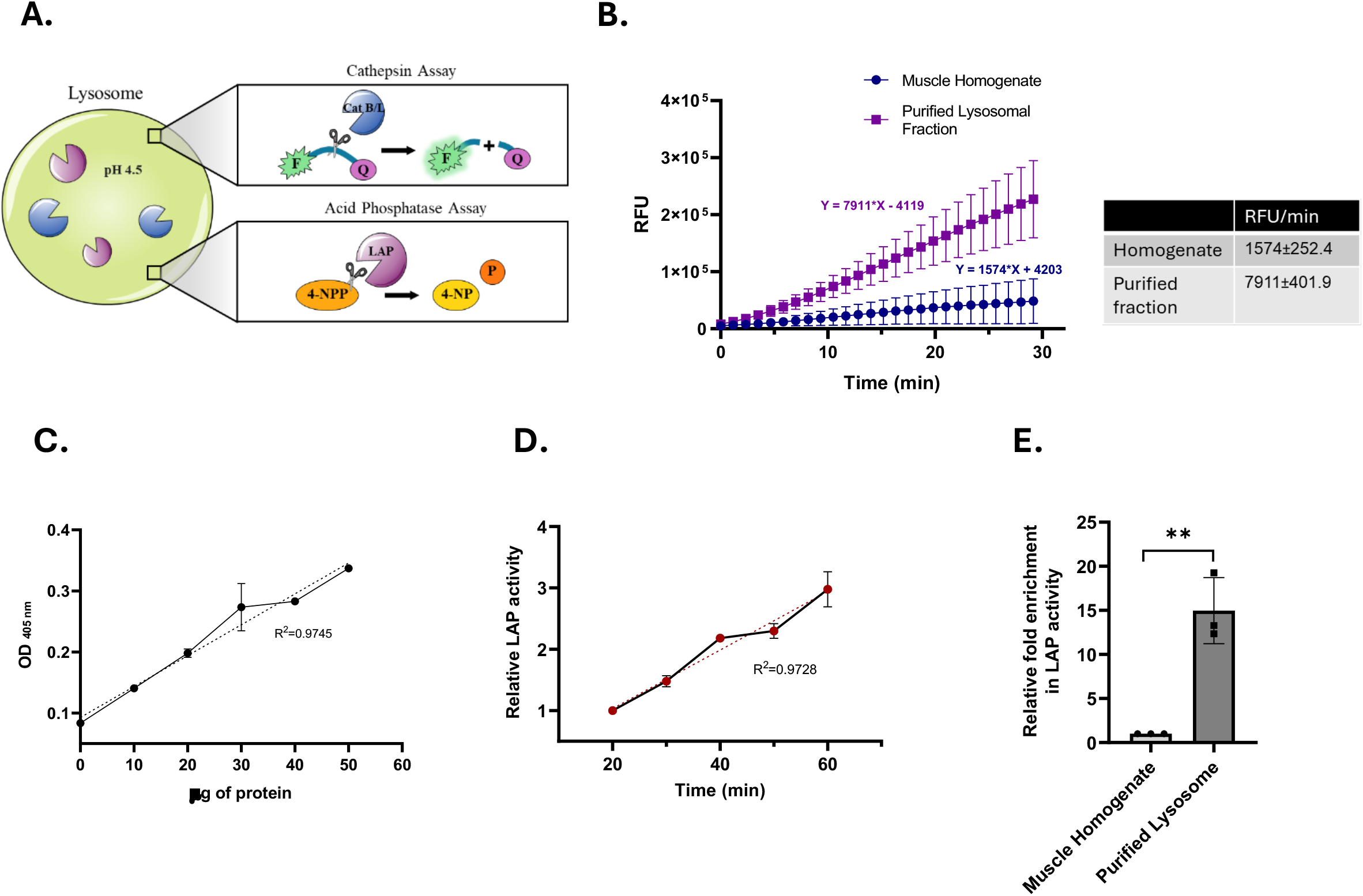
*A*: Schematic representation of the lysosomal Cathepsin B/L cleavage assay and acid phosphatase assay, illustrating a peptide substrate conjugated with a fluorophore and quencher. *B*: Relative fluorescence unit (RFU) of Cathepsin B/L cleavage assay in muscle homogenate and purified lysosomal fraction. *C*: Lysosomal acid phosphatase (LAP) activity in lysosome-enriched fractions measured by absorbance at 450 nm, following the reaction of varying amounts of purified fractions with the substrate. *D*: LAP activity in purified lysosomal fractions as a function of reaction time. *E*: Relative fold difference in LAP activity between muscle homogenate and purified lysosomal fractions. Values were determined based on OD_405 nm_ and are expressed as mean ± SD (n = 3; ^**^p < 0.005, unpaired t-test).

To further verify the integrity of our isolated lysosomes, we conducted a calcium kinetics assay using the FLUO-4 AM dye to mark calcium release, as lysosomes propagate calcium-based cellular signaling. The transient receptor potential mucolipin-1 (TRPML1) channel is a known calcium release channel at the site of the lysosome and is activated by the TRPML1-specific agent named mucolipin synthetic agonist 1 (ML-SA1). Following a basal kinetic read of approximately 20 seconds, which showed no differences in the FLUO-4 AM-based relative fluorescence, two isolated lysosomal samples were injected with DMSO or 10 μM of ML-SA1, followed by a 60-second kinetic read. ML-SA1 successfully released detectable calcium from the TRPML1 channel, which was not observed with DMSO treatment. This demonstrates not only the effectiveness of the release of calcium via TRPML1, but also the intactness of the isolated lysosomes, as intact lipid binding at the lysosomal membrane is required for detectable TRPML1 activity (27) (**Fig. 5**).

**Figure 5.**
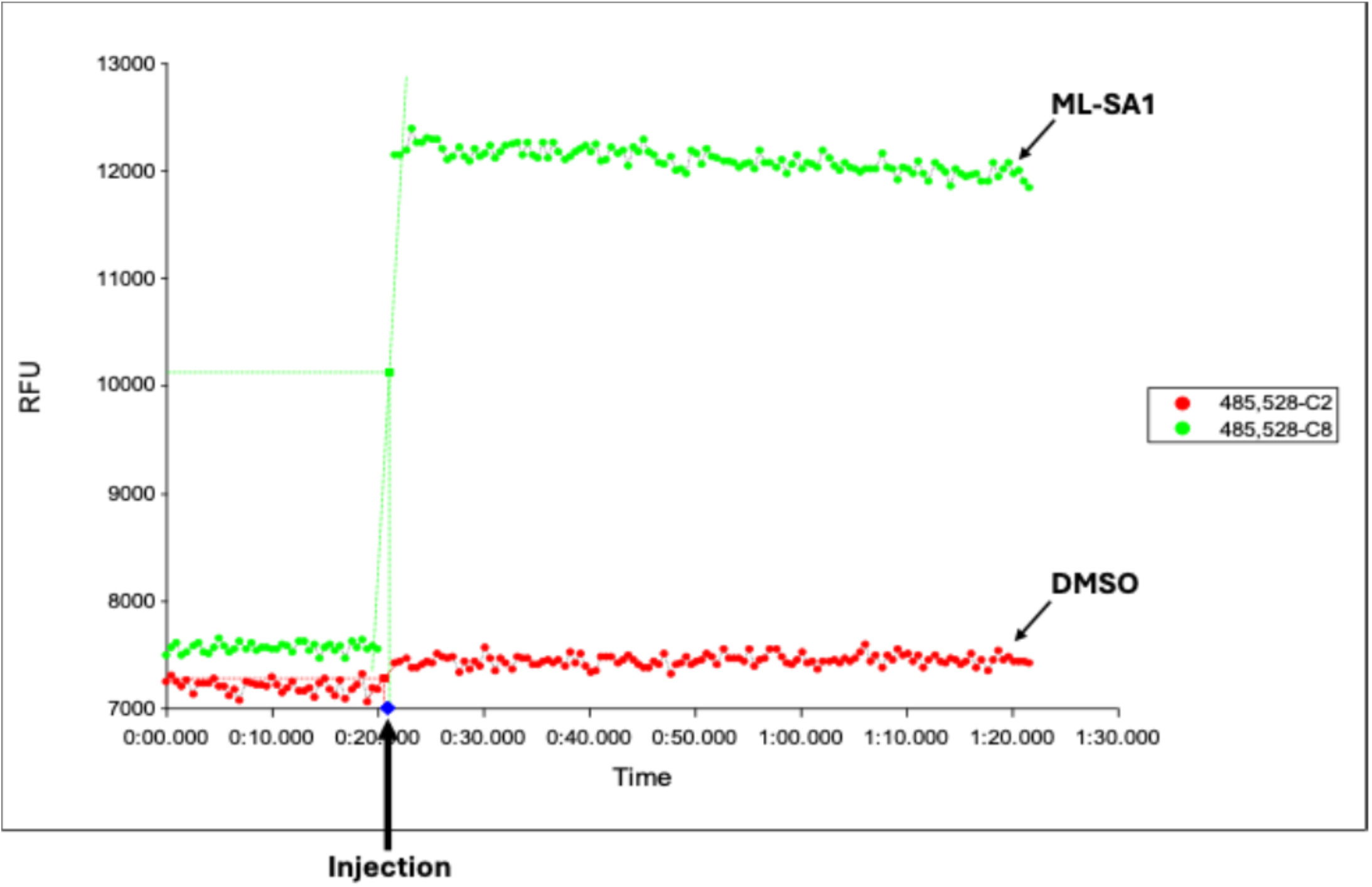
Representative tracings of FLUO-4 AM-stained lysosomes treated with DMSO (red tracing) or 10 uM ML-SA1 (green tracing) with an injection of either agent made after a 20-second basal kinetic read, followed by a 60-second post-injection kinetic read.

To evaluate the applicability of our lysosome isolation method in a physiologically relevant model, we applied this protocol to both young and aged skeletal muscle tissue. We then assessed lysosomal enzyme activity in these samples using the Cathepsin B activity assay to determine whether there are any age-related functional alterations. Notably, lysosomes isolated from aged muscle exhibited reduced enzymatic activity compared to those from young muscle (**Fig. 6A, 6B**).

**Figure 6.**
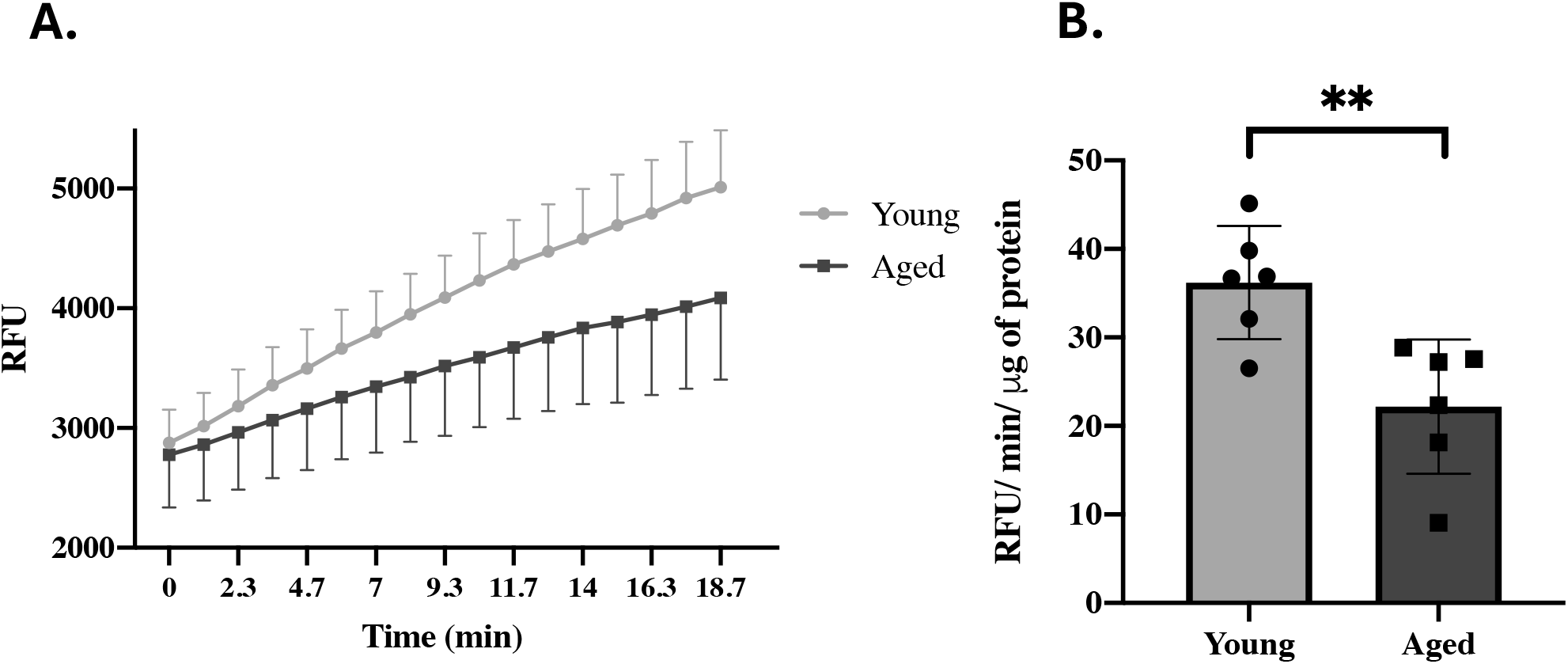
*A*: Relative fluorescence unit (RFU) of Cathepsin B/L cleavage assay in young and aged skeletal muscle purified lysosomal fractions. *B*: RFU per minute per μg of protein in young and aged skeletal muscle purified lysosomal fractions determined from the slope from *A*. Values were expressed as mean ± SD (n = 6; ^**^p < 0.005, unpaired t-test).

## Discussion

The isolation of lysosomes is crucial for analyzing lysosomal protein composition and function in skeletal muscle, thus providing insights into lysosome-mediated regulation and adaptation in health and disease. Lysosome isolation techniques have been established in tissues like the liver, brain, and kidney (12-14). However, isolating lysosomes from skeletal muscle presents unique challenges due to its distinct structural characteristics. The substantial presence of connective tissue, extracellular matrix, and structural proteins like collagen in skeletal muscle can interfere with tissue disruption and organelle separation during isolation (28). These factors necessitate the development of specialized protocols to effectively isolate lysosomes from skeletal muscle, ensuring sufficient yield and purity for downstream analyses. In this study, we developed a simple and efficient method to isolate lysosomes from skeletal muscle based on differential centrifugation without relying on external density gradients such as Percoll, Nycodenz, Iodixanol, or Cesium Chloride, commonly used in conventional subcellular fractionation protocols (13, 29, 30). To facilitate lysosome isolation, we utilized an isotonic buffer that maintains the osmolarity of the homogenate, minimizing organelle swelling and lysis during homogenization and subsequent isolation steps (31). It is essential to optimize the speed and time of tissue homogenization, depending on the mass of tissue used. During protocol development, we observed that elongated homogenization times and high speeds with relatively smaller mass (<0.5 g) caused the lysosomal rupture, leading to lysosomal enzyme release (**Supplementary Fig. 1**).

Compared to conventional protocols based on density gradient centrifugation that typically require 3–5 g of starting material and high-speed ultracentrifuges, our method offers a resource efficient alternative. Our protocol enables the isolation of lysosomes from as little as 0.5 g of muscle tissue, yielding approximately 100 –150 µg of purified lysosomes. Additionally, density gradient-based protocols have a higher risk of contamination with other organelles, such as mitochondria and peroxisomes, during layer separation. These methods also involve prolonged centrifugation at ultra-high speeds, which can result in lysosomal rupture and leakage of lysosomal enzymes, further compromising sample quality (32).

The approach of using Ca^2+^-mediated mitochondrial swelling in our protocol significantly enhanced the removal of mitochondrial contamination. While previous studies have employed Ca^2+^, they often required additional density gradients in the later stages to purify lysosomes. In contrast, our protocol eliminates the need for such gradients by optimizing the Ca^2+^ concentration to 8 mM with appropriate centrifugal speed and time. Conventional protocols have often involved injecting animals with Triton WR1339 and Dextran before lysosome isolation. These substances were used to modify lysosomal density, facilitating their separation from mitochondria and peroxisomes (12, 31). However, such administrations can lead to adverse effects in experimental animals. Further, the accumulation of these external molecules in lysosomes can significantly alter their natural biochemical properties and functions, leading to false interpretations of downstream functional analyses using isolated lysosomes. Our protocol avoids the use of such lysosomal density-altering substances and ensures that the lysosomes are isolated in their natural state. At the same time, recently two different high-yield purification methods have also been used to isolate lysosomes based on the usage of superparamagnetic nanoparticles (33) and lysosomal immunoprecipitation (34-36). These methods appear to be used mainly to isolate lysosomes from cultured cells, rather than from tissues. Another protocol for lysosome isolation involves affinity purification and immunoselection using antibodies against the vacuolar-type H(+) -ATPase (35). The protocol allows an efficient method to isolate lysosomes in ∼10 min from cultured cells and genetically modified tissues, compared to the protocol described here. However, this method relies on the use of large amounts of antibody-coated magnetic beads which can be costly and it requires genetic modification or the use of transgenic animals expressing TMEM192-3xHA, which limits the application of this protocol in wild-type tissues. A recent proteomic analysis also revealed that the overexpression of TMEM192 alters the expression of lysosomal membrane proteins and those involved in protein trafficking (37). Thus, under such circumstances, our traditional biochemical method provides a needed alternative to isolate lysosomes, preserving their natural protein composition.

The TEM images obtained following our isolation protocol confirmed the intactness of isolated lysosomes. It is essential to maintain the integrity of the lysosomal membrane, as any disruption to it could potentially lead to the release of enzymes, preventing the downstream lysosomal activity measurements. In addition, the stability and activity of lysosomal proteases rely on the acidic environment present within intact lysosomes (38). Further, any lysosomal rupture during the isolation could potentially reduce the final yield. The flow cytometry, TEM and calcium kinetic data proved that most of the lysosomes in the isolated fraction were intact, acidic and contained mucolipin channels.

Apart from analyzing lysosomal purity and intactness, measuring lysosomal function is critical to understanding their role in muscle homeostasis, metabolism, and disease. In this study, the use of acid phosphatase and cathepsin B assays has provided valuable insights into lysosomal function within skeletal muscle. Acid phosphatase activity, a key indicator of lysosomal enzyme function, has allowed us to assess the overall integrity and performance of the lysosomal system. Altered activity of this enzyme can indicate dysfunction, which is particularly relevant in muscle-related disorders where lysosomal degradation processes are impaired. Similarly, cathepsin B, a critical lysosomal protease involved in protein turnover and autophagy, has served as a more specific marker for proteolytic activity within the lysosome. By measuring cathepsin B activity, we can evaluate the efficiency of protein degradation, a process essential for maintaining muscle homeostasis, especially under conditions like exercise or injury. Furthermore, the isolated lysosome populations can be analyzed for mTORC1 activation and the presence of associated regulators such as Rag GTPases and Rheb to study lysosome-mediated nutrient signaling pathways in skeletal muscle. In case of lysophagy, certain markers of lysosomal damage and the recruitment of autophagy machinery will be detectable on isolated lysosomes. Specifically, the presence of Galectin-3, ubiquitin, p62, or LC3-II in lysosome-enriched fractions may serve as indirect indicators of lysophagy.

The results obtained from young and aged skeletal muscle support the use of this technique not only in healthy muscle but also in models of dysfunction, such as aging or disuse, thereby allowing the investigation of lysosome-related mechanisms in muscle pathophysiology in a variety of neurodegenerative or metabolic disorders in the future. In addition, this technique could also be utilized to assess lysosomal adaptations in response to chronic exercise training.

## Acknowledgements

This work was supported by grant funding from the Canadian Institutes of Health Research (CIHR) to D.A. Hood. Anastasiya Kuznyetsova was supported by a Canada Graduate Scholarship (CGS-M). Neushaw Moradi was supported by an Ontario Graduate Scholarship (OGS).

## Funding

Natural Science and Engineering Research Council (NSERC) and the Canadian Institutes of Health Research (CIHR)

## Conflict of Interest

None

## Notes

### Competing Interest Statement

The authors have declared no competing interest.

